# Case Study of High-Throughput Drug Screening and Remote Data Collection for SARS-CoV-2 Main Protease by Using Serial Femtosecond X-ray Crystallography

**DOI:** 10.1101/2021.11.28.468932

**Authors:** Omur Guven, Mehmet Gul, Esra Ayan, J. Austin Johnson, Baris Cakilkaya, Gozde Usta, Fatma Betul Ertem, Nurettin Tokay, Busra Yuksel, Oktay Gocenler, Cengizhan Buyukdag, Sabine Botha, Gihan Ketawala, Zhen Su, Brandon Hayes, Frederic Poitevin, Alexander Batyuk, Chun Hong Yoon, Christopher Kupitz, Serdar Durdagi, Raymond G. Sierra, Hasan DeMirci

**Author notes:** Equal contribution (Omur Guven, Mehmet Gul). Correspondence (H.D.).

## Abstract

Since early 2020, COVID-19 has grown to affect the lives of billions globally. A worldwide investigation has been ongoing for characterizing the virus and also for finding an effective drug and developing vaccines. As time has been of the essence, a crucial part of this research has been drug repurposing; therefore, confirmation of *in-silico* drug screening studies has been carried out for this purpose. Here we demonstrated the possibility of screening a variety of drugs efficiently by leveraging a high data collection rate of 120 images/second with the new low-noise, high dynamic range ePix10k2M Pixel Array Detector installed at the Macromolecular Femtosecond Crystallography (MFX) instrument at the Linac Coherent Light Source (LCLS). The X-ray Free-Electron Laser (XFEL) is used for remote high-throughput data collection for drug repurposing of the main protease (Mpro) of SARS-CoV-2 at ambient temperature with mitigated X-ray radiation damage. We obtained multiple structures soaked with 9 drug candidate molecules in two crystal forms. Although our drug binding attempts failed, we successfully established a high-throughput Serial Femtosecond X-ray crystallographic (SFX) data collection protocol.

## 1. Introduction

Emerging infectious diseases (EID) rapidly spread in a population [1]. Acquired immune deficiency syndrome (AIDS) [2], Lyme disease [3], H1N1 influenza [4], severe acute respiratory syndrome (SARS) [5], Zika, and Middle East respiratory syndrome (MERS) are among the recent outbreaks. Despite technological advances in healthcare, EIDs remain a very significant threat to global health [6]. In order to prevent and control the spread of such diseases, high-throughput approaches can play an essential role in drug repurposing and development.

SARS-CoV-2, like other coronaviruses, generates numerous polyproteins that must be processed into functional, unique proteins, such as nonstructural proteins required for viral replication [7]. SARS-CoV-2 genome encodes two proteases: papain-like cysteine protease (PLpro) and chymotrypsin-like cysteine protease, also called 3C-like protease (3CLpro) [8]. For the SARS-CoV-2 lifecycle, 3CLpro is necessary. This main protease 3CLpro (referred to as Mpro hereafter) is responsible for functional protein maturation and can be an attractive primary antiviral target [9]. Besides functional importance, targeting proteases, such as Mpro, is a common approach for combating viral infections because those proteins are highly conserved between species [10]. For instance, several Mpro drug targeting studies for SARS-CoV, MERS-CoV, and other viruses have been performed [11,12]; those efforts made the drug repurposing and design studies for SARS-CoV-2 Mpro possible due to a conserved active site and proteolytic activity [13]. Mpro is enzymatically active in the dimeric form, and its Cys-His catalytic dyad provides sequence-specific cleavage after Gln residues of polyproteins [10]. However, this conserved protein has no homologs or similar cleavage sites for proteases found in the human proteome, making it an ideal potential drug repurposing and screening target. Thus, drugs targeting Mpro are expected to have less or no side effects and toxicity while still acting as a broad-spectrum antiviral agent [14,15].

Drug-target complex structures determined at an atomic resolution are regularly used for drug repurposing [13]. X-ray Free-Electron Lasers (XFELs) are premier light sources for structure determination as they provide structures with mitigated radiation damage at ambient (near-physiological) temperatures. With high energy beams and 10-100 femtosecond (fs) pulse length, XFELs can help us to determine electronic states and molecular dynamics information that is invaluable for drug-repurposing studies. The comparatively brighter, short X-ray pulses, and the collection of diffraction data from exceedingly small (sub-micrometer) protein crystals make the use of hard XFELs attractive [16].

Previous structural studies conducted at the Linac Coherent Light Source (LCLS) facility have obtained two Mpro structures of SARS-CoV-2 with mitigated electron radiation damage, paving the way for potential drug repurposing studies for the treatment of COVID-19 [17]. High-throughput screening (HTS) is one of the methods used to discover small lead molecules and allowed development of new and higher quality drug candidates [18]. Various high-throughput screening studies with Nuclear Magnetic Resonance (NMR) spectroscopy are available in the literature. The most striking study was the high-throughput screening for the discovery of antiviral drugs against Ebola [19]. A similar study focuses on fragment screening for the DC-SIGN (Dendritic Cell-Specific Intercellular adhesion molecule-3-Grabbing Non-integrin) receptor. It is targeted by pathogens such as HIV, Ebola virus, and *Mycobacterium tuberculosis*. Within the scope of the study, five potential secondary target drug binding sites were identified [20]. In a recent study conducted at the Diamond Light Source, high-throughput crystallographic fragment screening was performed for SARS-CoV-2’s Nsp3 Mac 1 protein, which plays an essential role for viral replication. As a result, they identified numerous compounds that can be drug candidates against COVID-19 [21].

In this study, we report high-throughput X-ray diffraction data collection results for potential drug repurposing. Fifteen SARS-CoV-2 Mpro SFX structures were determined at the Macromolecular Femtosecond Crystallography (MFX) instrument of the LCLS at ambient temperature. XFELs provide micro-focused, ultrabright, and ultrafast X-ray pulses to probe micro- and nanocrystals in a serial fashion by obtaining individual snapshots and capturing Bragg reflection from exceedingly small crystals. It eliminates larger crystal growth requirements and simplifies optimization processes in batch mode, thus enabling it to work with multiple crystal forms and space groups. In a previous study, we obtained structures of SARS-CoV-2 Mpro at ambient temperature and verified the dynamic regions of the active site through the SFX approach [17]. This suggests immense potential by providing critical information for future high-throughput data collection for drug repurposing and computational modeling studies.

COVID-19 pandemic has shown us the importance of drug design/repurposing for infectious diseases and our workflow is an ideal suited for future drug repurposing and screening studies. The method is fast, effective, and physiologically relevant since the structures are obtained at ambient temperature.

## 2. Materials and Methods

### 2.1. Gene construct design, protein expression and purification

Two Mpro gene constructs were designed, expressed, and purified as stated in [17]. Two Mpro gene constructs were ordered from Genscript, USA in a pET28a(+) *Escherichia coli* expression vector. One of our constructs had an N-terminus hexahistidine-tag with a native Mpro cleavage site and a C-terminal PreScission™ cut site. After the protease cuts, Mpro is left with native N- and C-terminals; this construct is referred to as the “native construct”. The other construct had an N-terminal thrombin cut site that leaves an additional 4 residues upon cleavage (GSHM); hence, it is called the “modified construct”. Bacterial transformations were performed on a chemically competent *Escherichia coli* BL21 Rosetta-2 strain for both constructs. Transformed cells were grown in 12 L LB-Miller media supplemented with 50 μl/ml kanamycin and 35 μl/ml chloramphenicol antibiotics at 37°C. When the OD600 reached ~1.0, isopropyl β-D-1-thiogalactopyranoside (IPTG) induction was done with a final concentration of 0.4 mM IPTG for protein production. Cells were harvested at 3500 rpm for 20 minutes at 4°C by using Beckman Allegra 15R desktop centrifuge. Supernatant was discarded and cell pellets were stored at −80°C until protein purification. The pellet was dissolved in a lysis buffer containing 50 mM Tris-HCl (pH 7.5), 300 mM NaCl and 5% v/v glycerol supplemented with 0.01% Triton X-100 by sonicating the solution (Branson W250 sonifier, USA). Cell lysate was ultracentrifuged (The Beckman Optima™ L-80 XP Ultracentrifuge) at 40000 rpm for 30 min at 4°C with a Ti45 rotor (Beckman, USA). Mpro-containing supernatant was applied to an immobilized metal affinity chromatography (IMAC) column by using a Ni-NTA agarose resin (QIAGEN, USA) with a constant flow rate of 2 ml/min. The column was washed with a loading buffer that contains 20 mM Tris-HAc (pH 7.5), 150 mM NaCl, and 5 mM Imidazole to remove non-specific bindings. To elute the Mpro, 20 mM Tris-HAc (pH 7.5), 150 mM NaCl, 250 mM Imidazole elution buffer was applied, and 35 ml of elute was collected. Elute was dialyzed against 20 mM Tris-HAc (pH 7.5), 150 mM NaCl overnight in a 3 kDa cut-off dialysis membrane to remove the excess Imidazole. For the native construct, 3C protease (Prescission protease, GenScript, USA) was used in a 1:100 stoichiometric molar ratio to cleave the hexahistidine-tag. For the modified Mpro construct, thrombin protease (Sigma, USA) was used in a 1:100 stoichiometric molar ratio to cut off the hexahistidine from the protein. After overnight incubation with proteases at 4°C, elute was applied to reverse Ni-NTA chromatography to purify the untagged protein from the remaining part of the affinity tag in the solution. Mpro proteins were concentrated to 25 mg/ml by Millipore’s ultrafiltration columns and a final concentration of 1 mM DTT was added and stored at −80°C.

### 2.2. Crystallization of protein with drug by using soaking method

For protein crystallization, high-throughput sitting drop microbatch under paraffin oil screening was performed under paraffin oil. Purified protein with 25 mg/ml concentration was mixed with sparse matrix crystallization screening conditions in a 1:1 volumetric ratio by using 72 well Terasaki crystallization plates (Greiner-Bio, Germany). The mixture was sealed with 100% paraffin oil to slow down the vapor diffusion (Tekkim Kimya, Turkey) and crystallization plates are stored at ambient temperature. The best crystals for the native construct were obtained from Pact Premier™ crystallization screen 1 condition #39 (0.1 M MMT pH 6.0, 25% w/v PEG 1500) (Molecular Dimensions). For the modified construct, crystals were obtained from Crystal Screen™ crystallization screen condition #22 (0.2 M Sodium acetate trihydrate, 0.1 M Tris pH 8.5, 30 % w/v PEG 4000) (Hampton Research). Successful crystal conditions were replicated at a larger scale where 10 ml of the protein sample was mixed with 10 ml of the referred comercial crystallization conditions. After crystals of size 1-5 μm × 5-10 μm × 10-20 μm reached, they were soaked with 100 μg/mL of drug solution by mixing the drug solutions into protein crystal solution for both constructs. Promising drug candidates were chosen by intensive literature search: Lopinavir [22], Rosuvastatin and Atorvastatin [23], Adefovir [24], Umifenovir [25], Empagliflozin [26], Ebselen [17], Montelukast [27,28].

### 2.3. Transport of microcrystals

Crystals were transferred into screw-top plastic cryovials (Wuxi NEST biotechnology, China cat#607001). The cryovials were loosely wrapped with Kimwipes (Kimberly-Clark, USA) to absorb mechanical shocks during the transport from Istanbul, Turkey to Menlo Park, CA, USA. These cryovials were placed inside larger screw-top glass vials. The glass vials were covered with extra layers of cotton (Ipek, Turkey) and placed within a Ziploc™ bag (SC Johnson, USA) for further mechanical shock absorption and providing additional thermal insulation. For further thermal insulation, the Ziploc™ bags were placed in a styrofoam box filled with ~1 kg of cotton during transportation. The styrofoam box was tightly closed and covered with an additional 1 cm thick layer of cotton, duct-taped all around to prevent thermal fluctuations during international transport.

### 2.4. Injection of microcrystals

The sample reservoir, which has a volume of 1.6 mL, was filled with soaked crystal slurries in their unmodified mother liquor. To utilize the electrospinning method, a standard Microfluidic Electrokinetic Sample Holder (MESH) was used for sample injection, allowing us to collect full datasets using a reduced amount of sample [29,30]. The sample capillary consists of a 200 μm ID × 360 μm OD × 1.0 m long capillary made of fused silica. The voltage applied to the sample fluid was 2500-3000 V, and the counter electrode was grounded. The sample flow rate was between 2.5 and 8 μl/min in general.

### 2.5. Data collection at LCLS

The SFX experiments with microcrystals were performed at the MFX instrument at the LCLS (beamtime ID: mfx17318) at the SLAC National Accelerator Laboratory (Menlo Park, CA). An X-ray beam with a vertically polarized pulse of 30-40 fs duration was focused with refractive beryllium compound lenses with a ~6×6 μm full width at half maximum beam size. The total pulse energy was 0.8 mJ, with 9.8 keV (1.25 Å) photon energy, 10^12^ photons per pulse flux and a 120 Hz repetition rate. *OM monitor* [31] and *PSOCAKE* [32, 33] were used to determine the initial diffraction geometry, monitor the crystal hit rates, and analyze the gain switching modes of the epix10k 2-megapixel (ePix10k2M) detector [34]. The detector frames were collected continuously from Mpro microcrystals, and the total number of frames collected per datasets ranged from 192352 to 1465292 (Table 1). Total run times were between 36min and 5h 20min with the effective run times ranging from 36 mins to 4hrs 20mins (Table 1). The ePix10k2M detector was used in dynamic gain mode activated. During the data collection the MESH injector system had no blockages. The total number of frames with more than 30 Bragg reflections and a signal-to-noise ratio larger than 4.5 were considered as a hit and their numbers were ranged from 5274 to 45521 per dataset (Table 1). The distance for the detector was 118 mm that has a 2.1 Å achievable resolution on the edges and 1.64 Å in the corners. All the relevant information for the data collection process is further summarized in Table 1.

**Table 1.** Serial femtosecond X-ray data collection statistics.

| Sample ID | PDB ID | Total Run Time | Effective Run Time | Runs | Run Number | Total number frames collected | Total number frames with more than 30 Bragg reflections and a signal to noise ratio > 4.5 | Hit rate (%) | Total number of crystal lattices indexed to the appropriate unit cell | Total number of crystal lattices merged into the final SFX dataset |
| --- | --- | --- | --- | --- | --- | --- | --- | --- | --- | --- |
| Mpro3 | 7VJY | 5h 20m | 4h 20m | 20-80 | 60 | 1465292 | 279428 | 19.07 | 73605 | 68314 |
| Mpro4 | 7VK3 | 3h 28m | 2h 59m | 82-115 | 33 | 1121297 | 182774 | 16.3 | 33384 | 67918 |
| Mpro5 | 7CWB | 2h 59m | 2h 52m | 118-144 | 26 | 1163413 | 208839 | 17.95 | 168655 | 160510 |
| Mpro6 | 7VK4 | 2h 18m | 2h 18m | 235-257 | 22 | 790492 | 229422 | 29.02 | 315158 | 313250 |
| Mpro7 | 7VK8 | 1h 25m | 45m | 176-188 | 12 | 327156 | 2078 | 0.63 | 1177 | 947 |
| Mpro8 | N/A | N/A | N/A | N/A | N/A | N/A | 13 | N/A | N/A | N/A |
| Mpro9 | 7VK1 | 3h 04m | 3h 04m | 145-174 | 29 | 1148580 | 51112 | 4.45 | 33384 | 30732 |
| Mpro10 | 7VK0 | 2h 11m | 2h 11m | 264-291 | 27 | 858742 | 93235 | 10.86 | 116315 | 115577 |
| Mpro11 | 7VJZ | 2h 07m | 1h 57m | 189-213 | 24 | 669237 | 132447 | 19.79 | 70272 | 65579 |
| Mpro12 | 7VK5 | 1h 30m | 1h 30m | 292-310 | 18 | 609036 | 80383 | 13.19 | 120147 | 119430 |
| Mpro13 | 7VK2 | 2h 47m | 2h 47 | 216-234 | 18 | 873323 | 180625 | 20.68 | 88551 | 81563 |
| Mpro14 | 7VJW | 1h 12m | 1h 12m | 312-234 | 7 | 192352 | 15899 | 8.26 | 22470 | 21278 |
| Mpro15 | N/A | N/A | N/A | N/A | N/A | N/A | 0 | N/A | N/A | N/A |
| Mpro16 | 7CWC | 1h 53m | 1h 53m | 320-340 | 20 | 686808 | 214355 | 31.21 | 157976 | 156512 |
| Mpro17 | 7VK7 | 1h 28m | 1h 28m | 341-357 | 16 | 439830 | 86490 | 19.66 | 174120 | 173796 |
| Mpro18 | 7VJX | 1h 22m | 1h 22m | 358-372 | 15 | 466014 | 111815 | 23.99 | 170214 | 169695 |
| Mpro19 | 7VK6 | 36m | 36m | 373-375 | 3 | 326781 | 78455 | 24.01 | 111145 | 107895 |

### 2.6. Data processing

Data processing details were performed as described in the article by Durdagi *et al*. [17]. The hit findings and detector correction were performed through the *CHEETAH* software [35]. All data sets were classified according to the hit finding parameters created using *peakfinder8* [35]. The crystal hits were indexed through *CrystFEL* version 0.9.0 [36] software package and specific indexing algorithms. Indexed reflections integrated and merged through *PARTIALATOR* [37] were obtained as a complete reflection intensity list from *CrystFEL* [36]. These intensities were scaled and trimmed through the *TRUNCATE* [38] program. The indexed patterns were merged as the final dataset.

### 2.7. Structure determination

Structure determination was performed as described in the article by Durdagi *et al*. [17]. The determination of Mpro crystal forms was accomplished by the automated molecular replacement (MR) program *PHASER* [39]. For MR phasing of the structures in the space group C121, we used the coordinates of PDB ID: 7CWB [17] as our initial search model. For the structures in the space group P2_1_2_1_2_1_, we used the coordinates of PDB ID: 7CWC [17] as our initial MR search model. We performed our initial rigid-body refinements through the *PHENIX* [40] software package. The positions of altered side chains were defined by completing simulated-annealing refinement followed by individual coordinates and TLS parameter refinement. The water molecules and strong difference density positions of the structures were checked using the *COOT* [41] software. Structural alignments and all figures were generated using the *PyMOL* [42] software. The statistics for the structure determination are summarized in Table 2.

**Table 2.**
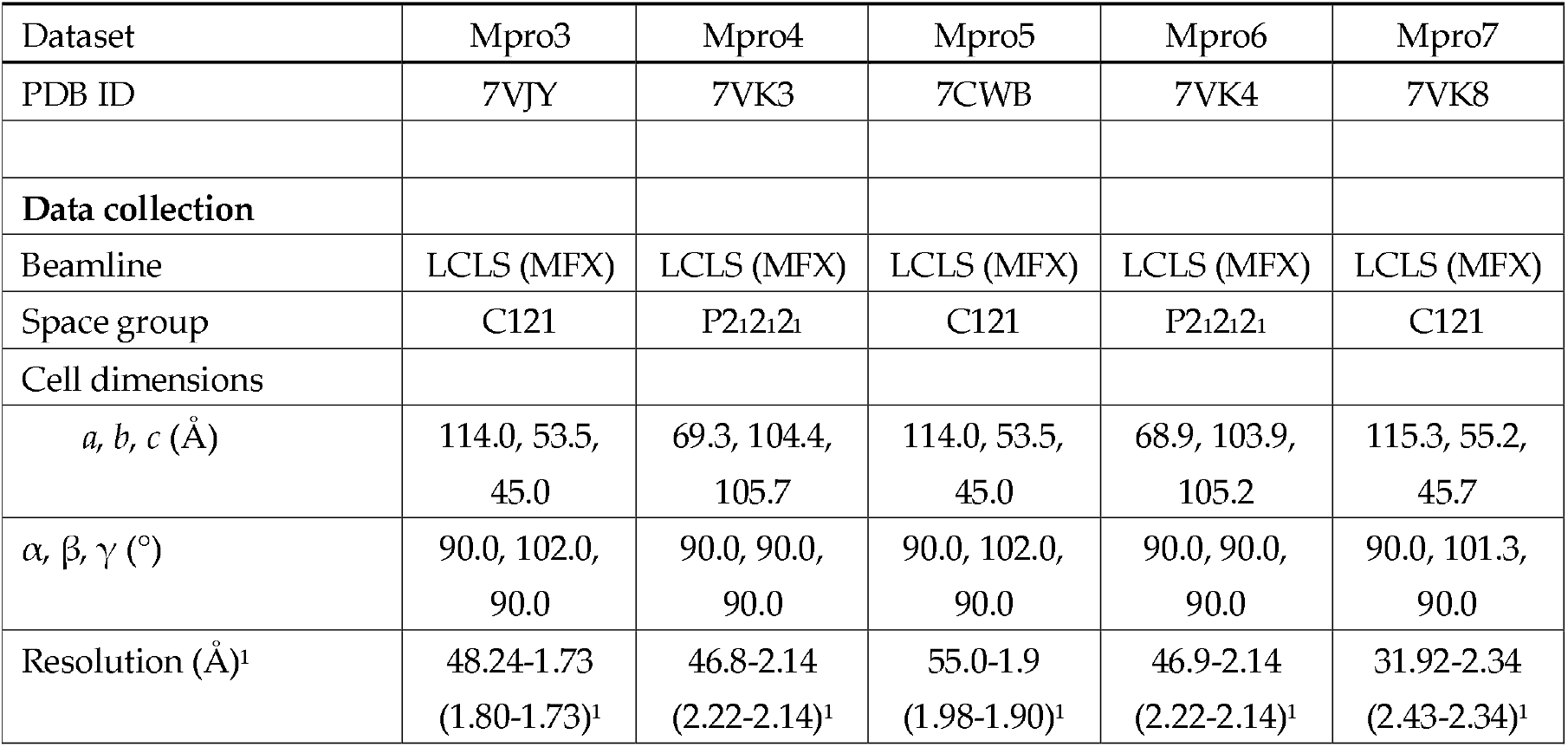
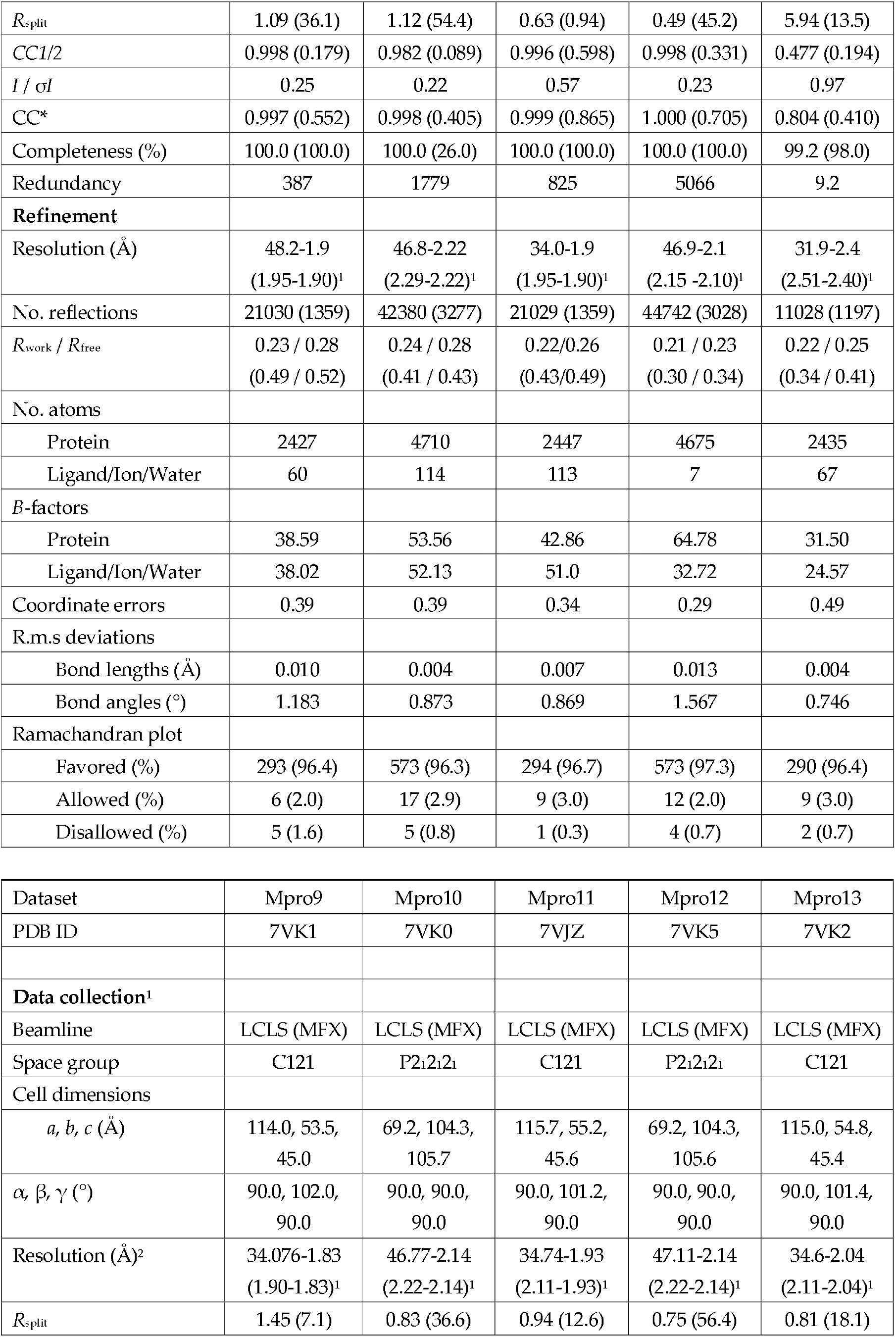
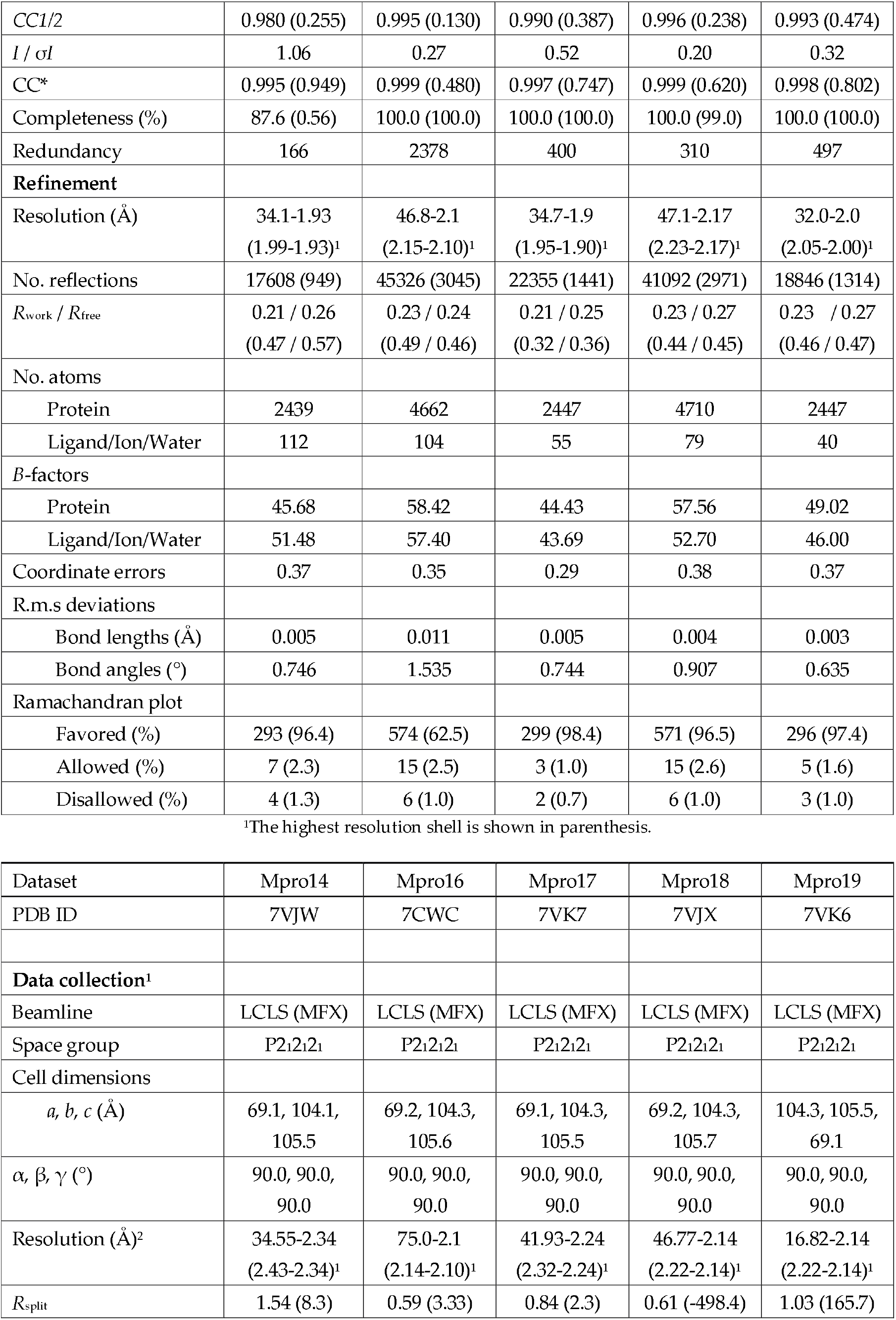
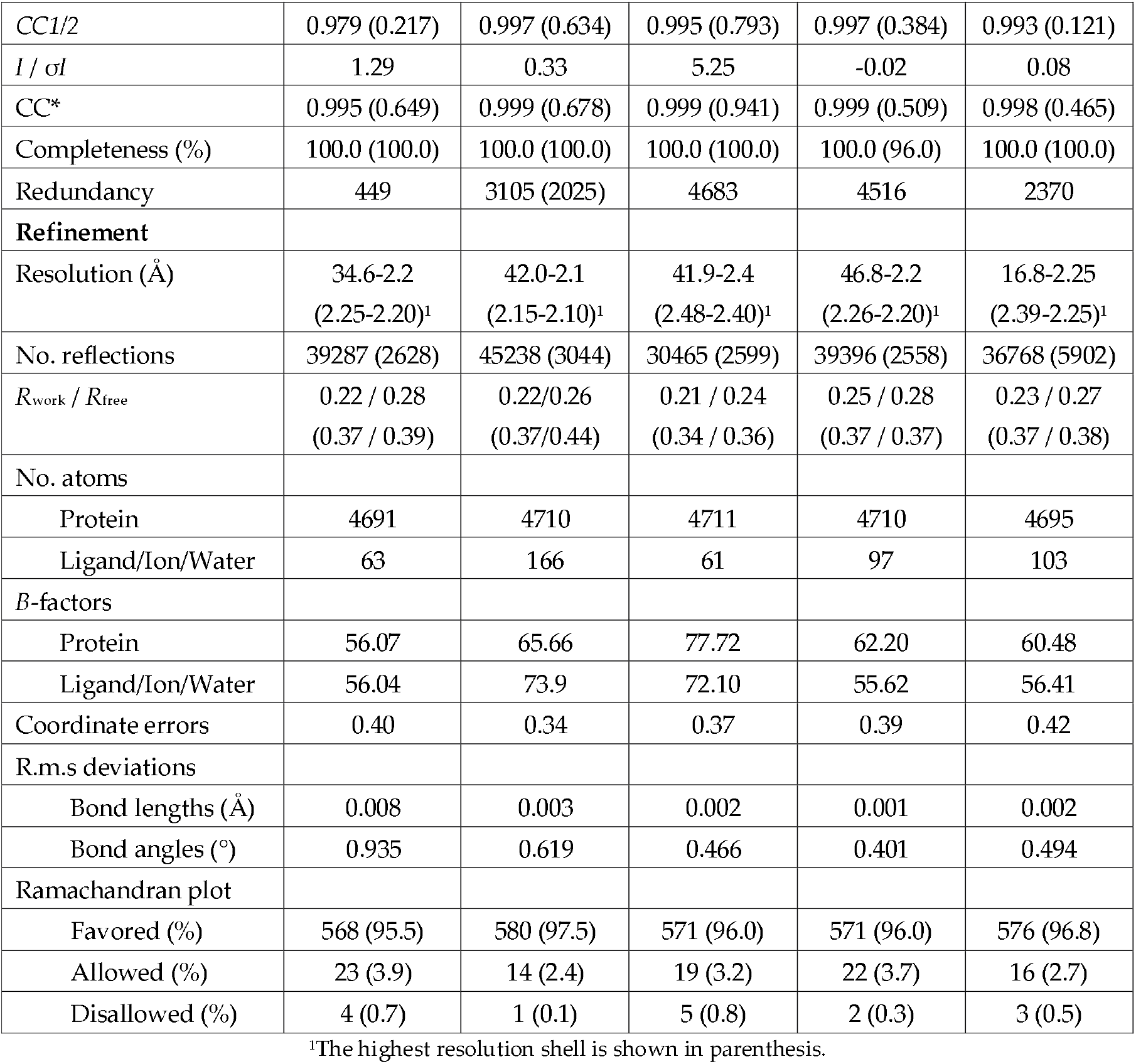
Data collection and refinement statistics for X-ray crystallography.

| Dataset | Mpro3 | Mpro4 | Mpro5 | Mpro6 | Mpro7 |
| --- | --- | --- | --- | --- | --- |
| PDB ID | 7VJY | 7VK3 | 7CWB | 7VK4 | 7VK8 |
| **Data collection** |  |  |  |  |  |
| Beamline | LCLS (MFX) | LCLS (MFX) | LCLS (MFX) | LCLS (MFX) | LCLS (MFX) |
| Space group | C121 | P2<sub>1</sub>2<sub>1</sub>2<sub>1</sub> | C121 | P2<sub>1</sub>2<sub>1</sub>2<sub>1</sub> | C121 |
| Cell dimensions |  |  |  |  |  |
| *a*, *b*, *c* (Å) | 114.0, 53.5,<br>45.0 | 69.3, 104.4,<br>105.7 | 114.0, 53.5,<br>45.0 | 68.9, 103.9,<br>105.2 | 115.3, 55.2,<br>45.7 |
| $\alpha$, $\beta$, $\gamma$ (°) | 90.0, 102.0,<br>90.0 | 90.0, 90.0,<br>90.0 | 90.0, 102.0,<br>90.0 | 90.0, 90.0,<br>90.0 | 90.0, 101.3,<br>90.0 |
| Resolution (Å)<sup>1</sup> | 48.24-1.73<br>(1.80-1.73)<sup>1</sup> | 46.8-2.14<br>(2.22-2.14)<sup>1</sup> | 55.0-1.9<br>(1.98-1.90)<sup>1</sup> | 46.9-2.14<br>(2.22-2.14)<sup>1</sup> | 31.92-2.34<br>(2.43-2.34)<sup>1</sup> |
| $R_{\text{split}}$ | 1.09 (36.1) | 1.12 (54.4) | 0.63 (0.94) | 0.49 (45.2) | 5.94 (13.5) |
| $CC1/2$ | 0.998 (0.179) | 0.982 (0.089) | 0.996 (0.598) | 0.998 (0.331) | 0.477 (0.194) |
| $I / \sigma I$ | 0.25 | 0.22 | 0.57 | 0.23 | 0.97 |
| $CC^*$ | 0.997 (0.552) | 0.998 (0.405) | 0.999 (0.865) | 1.000 (0.705) | 0.804 (0.410) |
| Completeness (%) | 100.0 (100.0) | 100.0 (26.0) | 100.0 (100.0) | 100.0 (100.0) | 99.2 (98.0) |
| Redundancy | 387 | 1779 | 825 | 5066 | 9.2 |
| **Refinement** |  |  |  |  |  |
| Resolution (Å) | 48.2-1.9<br>(1.95-1.90)<sup>1</sup> | 46.8-2.22<br>(2.29-2.22)<sup>1</sup> | 34.0-1.9<br>(1.95-1.90)<sup>1</sup> | 46.9-2.1<br>(2.15 -2.10)<sup>1</sup> | 31.9-2.4<br>(2.51-2.40)<sup>1</sup> |
| No. reflections | 21030 (1359) | 42380 (3277) | 21029 (1359) | 44742 (3028) | 11028 (1197) |
| $R_{\text{work}} / R_{\text{free}}$ | 0.23 / 0.28<br>(0.49 / 0.52) | 0.24 / 0.28<br>(0.41 / 0.43) | 0.22/0.26<br>(0.43/0.49) | 0.21 / 0.23<br>(0.30 / 0.34) | 0.22 / 0.25<br>(0.34 / 0.41) |
| No. atoms |  |  |  |  |  |
| Protein | 2427 | 4710 | 2447 | 4675 | 2435 |
| Ligand/Ion/Water | 60 | 114 | 113 | 7 | 67 |
| $B$-factors | | | | | |
| Protein | 38.59 | 53.56 | 42.86 | 64.78 | 31.50 |
| Ligand/Ion/Water | 38.02 | 52.13 | 51.0 | 32.72 | 24.57 |
| Coordinate errors | 0.39 | 0.39 | 0.34 | 0.29 | 0.49 |
| R.m.s deviations |  |  |  |  |  |
| Bond lengths (Å) | 0.010 | 0.004 | 0.007 | 0.013 | 0.004 |
| Bond angles (°) | 1.183 | 0.873 | 0.869 | 1.567 | 0.746 |
| Ramachandran plot |  |  |  |  |  |
| Favored (%) | 293 (96.4) | 573 (96.3) | 294 (96.7) | 573 (97.3) | 290 (96.4) |
| Allowed (%) | 6 (2.0) | 17 (2.9) | 9 (3.0) | 12 (2.0) | 9 (3.0) |
| Disallowed (%) | 5 (1.6) | 5 (0.8) | 1 (0.3) | 4 (0.7) | 2 (0.7) |
| Dataset | Mpro9 | Mpro10 | Mpro11 | Mpro12 | Mpro13 |
| PDB ID | 7VK1 | 7VK0 | 7VJZ | 7VK5 | 7VK2 |
| **Data collection<sup>1</sup>** |  |  |  |  |  |
| Beamline | LCLS (MFX) | LCLS (MFX) | LCLS (MFX) | LCLS (MFX) | LCLS (MFX) |
| Space group | C121 | P2<sub>1</sub>2<sub>1</sub>2<sub>1</sub> | C121 | P2<sub>1</sub>2<sub>1</sub>2<sub>1</sub> | C121 |
| Cell dimensions |  |  |  |  |  |
| $a, b, c$ (Å) | 114.0, 53.5,<br>45.0 | 69.2, 104.3,<br>105.7 | 115.7, 55.2,<br>45.6 | 69.2, 104.3,<br>105.6 | 115.0, 54.8,<br>45.4 |
| $\alpha, \beta, \gamma$ (°) | 90.0, 102.0,<br>90.0 | 90.0, 90.0,<br>90.0 | 90.0, 101.2,<br>90.0 | 90.0, 90.0,<br>90.0 | 90.0, 101.4,<br>90.0 |
| Resolution (Å)<sup>2</sup> | 34.076-1.83<br>(1.90-1.83)<sup>1</sup> | 46.77-2.14<br>(2.22-2.14)<sup>1</sup> | 34.74-1.93<br>(2.11-1.93)<sup>1</sup> | 47.11-2.14<br>(2.22-2.14)<sup>1</sup> | 34.6-2.04<br>(2.11-2.04)<sup>1</sup> |
| $R_{\text{split}}$ | 1.45 (7.1) | 0.83 (36.6) | 0.94 (12.6) | 0.75 (56.4) | 0.81 (18.1) |
| *CC1/2* | 0.980 (0.255) | 0.995 (0.130) | 0.990 (0.387) | 0.996 (0.238) | 0.993 (0.474) |
| *I* / $\sigma I$ | 1.06 | 0.27 | 0.52 | 0.20 | 0.32 |
| *CC\** | 0.995 (0.949) | 0.999 (0.480) | 0.997 (0.747) | 0.999 (0.620) | 0.998 (0.802) |
| Completeness (%) | 87.6 (0.56) | 100.0 (100.0) | 100.0 (100.0) | 100.0 (99.0) | 100.0 (100.0) |
| Redundancy | 166 | 2378 | 400 | 310 | 497 |
| **Refinement** |  |  |  |  |  |
| Resolution (Å) | 34.1-1.93<br>(1.99-1.93)<sup>1</sup> | 46.8-2.1<br>(2.15-2.10)<sup>1</sup> | 34.7-1.9<br>(1.95-1.90)<sup>1</sup> | 47.1-2.17<br>(2.23-2.17)<sup>1</sup> | 32.0-2.0<br>(2.05-2.00)<sup>1</sup> |
| No. reflections | 17608 (949) | 45326 (3045) | 22355 (1441) | 41092 (2971) | 18846 (1314) |
| *R*<sub>work</sub> / *R*<sub>free</sub> | 0.21 / 0.26<br>(0.47 / 0.57) | 0.23 / 0.24<br>(0.49 / 0.46) | 0.21 / 0.25<br>(0.32 / 0.36) | 0.23 / 0.27<br>(0.44 / 0.45) | 0.23 / 0.27<br>(0.46 / 0.47) |
| No. atoms |  |  |  |  |  |
| Protein | 2439 | 4662 | 2447 | 4710 | 2447 |
| Ligand/Ion/Water | 112 | 104 | 55 | 79 | 40 |
| *B*-factors |  |  |  |  |  |
| Protein | 45.68 | 58.42 | 44.43 | 57.56 | 49.02 |
| Ligand/Ion/Water | 51.48 | 57.40 | 43.69 | 52.70 | 46.00 |
| Coordinate errors | 0.37 | 0.35 | 0.29 | 0.38 | 0.37 |
| R.m.s deviations |  |  |  |  |  |
| Bond lengths (Å) | 0.005 | 0.011 | 0.005 | 0.004 | 0.003 |
| Bond angles (°) | 0.746 | 1.535 | 0.744 | 0.907 | 0.635 |
| **Ramachandran plot** |  |  |  |  |  |
| Favored (%) | 293 (96.4) | 574 (62.5) | 299 (98.4) | 571 (96.5) | 296 (97.4) |
| Allowed (%) | 7 (2.3) | 15 (2.5) | 3 (1.0) | 15 (2.6) | 5 (1.6) |
| Disallowed (%) | 4 (1.3) | 6 (1.0) | 2 (0.7) | 6 (1.0) | 3 (1.0) |

<sup>1</sup>The highest resolution shell is shown in parenthesis.

| Dataset | Mpro14 | Mpro16 | Mpro17 | Mpro18 | Mpro19 |
| --- | --- | --- | --- | --- | --- |
| PDB ID | 7VJW | 7CWC | 7VK7 | 7VJX | 7VK6 |
| **Data collection<sup>1</sup>** |  |  |  |  |  |
| Beamline | LCLS (MFX) | LCLS (MFX) | LCLS (MFX) | LCLS (MFX) | LCLS (MFX) |
| Space group | P2<sub>1</sub>2<sub>1</sub>2<sub>1</sub> | P2<sub>1</sub>2<sub>1</sub>2<sub>1</sub> | P2<sub>1</sub>2<sub>1</sub>2<sub>1</sub> | P2<sub>1</sub>2<sub>1</sub>2<sub>1</sub> | P2<sub>1</sub>2<sub>1</sub>2<sub>1</sub> |
| Cell dimensions |  |  |  |  |  |
| *a*, *b*, *c* (Å) | 69.1, 104.1,<br>105.5 | 69.2, 104.3,<br>105.6 | 69.1, 104.3,<br>105.5 | 69.2, 104.3,<br>105.7 | 104.3, 105.5,<br>69.1 |
| $\alpha$, $\beta$, $\gamma$ (°) | 90.0, 90.0,<br>90.0 | 90.0, 90.0,<br>90.0 | 90.0, 90.0,<br>90.0 | 90.0, 90.0,<br>90.0 | 90.0, 90.0,<br>90.0 |
| Resolution (Å)<sup>2</sup> | 34.55-2.34<br>(2.43-2.34)<sup>1</sup> | 75.0-2.1<br>(2.14-2.10)<sup>1</sup> | 41.93-2.24<br>(2.32-2.24)<sup>1</sup> | 46.77-2.14<br>(2.22-2.14)<sup>1</sup> | 16.82-2.14<br>(2.22-2.14)<sup>1</sup> |
| *R*<sub>split</sub> | 1.54 (8.3) | 0.59 (3.33) | 0.84 (2.3) | 0.61 (-498.4) | 1.03 (165.7) |
| *CC1/2* | 0.979 (0.217) | 0.997 (0.634) | 0.995 (0.793) | 0.997 (0.384) | 0.993 (0.121) |
| *I* / $\sigma I$ | 1.29 | 0.33 | 5.25 | -0.02 | 0.08 |
| *CC\** | 0.995 (0.649) | 0.999 (0.678) | 0.999 (0.941) | 0.999 (0.509) | 0.998 (0.465) |
| Completeness (%) | 100.0 (100.0) | 100.0 (100.0) | 100.0 (100.0) | 100.0 (96.0) | 100.0 (100.0) |
| Redundancy | 449 | 3105 (2025) | 4683 | 4516 | 2370 |
| **Refinement** |  |  |  |  |  |
| Resolution (Å) | 34.6-2.2<br>(2.25-2.20)<sup>1</sup> | 42.0-2.1<br>(2.15-2.10)<sup>1</sup> | 41.9-2.4<br>(2.48-2.40)<sup>1</sup> | 46.8-2.2<br>(2.26-2.20)<sup>1</sup> | 16.8-2.25<br>(2.39-2.25)<sup>1</sup> |
| No. reflections | 39287 (2628) | 45238 (3044) | 30465 (2599) | 39396 (2558) | 36768 (5902) |
| *R*<sub>work</sub> / *R*<sub>free</sub> | 0.22 / 0.28<br>(0.37 / 0.39) | 0.22/0.26<br>(0.37/0.44) | 0.21 / 0.24<br>(0.34 / 0.36) | 0.25 / 0.28<br>(0.37 / 0.37) | 0.23 / 0.27<br>(0.37 / 0.38) |
| No. atoms |  |  |  |  |  |
| Protein | 4691 | 4710 | 4711 | 4710 | 4695 |
| Ligand/Ion/Water | 63 | 166 | 61 | 97 | 103 |
| *B*-factors |  |  |  |  |  |
| Protein | 56.07 | 65.66 | 77.72 | 62.20 | 60.48 |
| Ligand/Ion/Water | 56.04 | 73.9 | 72.10 | 55.62 | 56.41 |
| Coordinate errors | 0.40 | 0.34 | 0.37 | 0.39 | 0.42 |
| R.m.s deviations |  |  |  |  |  |
| Bond lengths (Å) | 0.008 | 0.003 | 0.002 | 0.001 | 0.002 |
| Bond angles (°) | 0.935 | 0.619 | 0.466 | 0.401 | 0.494 |
| Ramachandran plot |  |  |  |  |  |
| Favored (%) | 568 (95.5) | 580 (97.5) | 571 (96.0) | 571 (96.0) | 576 (96.8) |
| Allowed (%) | 23 (3.9) | 14 (2.4) | 19 (3.2) | 22 (3.7) | 16 (2.7) |
| Disallowed (%) | 4 (0.7) | 1 (0.1) | 5 (0.8) | 2 (0.3) | 3 (0.5) |

<sup>1</sup>The highest resolution shell is shown in parenthesis.

## 3. Results

### 3.1. Serial crystallography based faster high-throughput drug screening at an XFEL

During our LCLS beamtime (ID: mfx17318), the new-generation detector ePix10k2M was employed in dynamic gain mode. The new hybrid pixel detector ePix10k2M runs at repetition rates of 120 Hz up to 1 kHz, which allows much faster data acquisition rates compared to its predecessors [43]. During data collection, we did not experience any blockages with the MESH injector system. As a result, a total of 11,304,510 detector frames were collected in 1 day and 07h 08m with the new detector from Mpro microcrystals. Fifteen datasets were collected from 15 crystal soaks of Mpro with various potential drug candidates. The Mpro03 datasets were collected in 5h20m as it was an early dataset; however, the Mpro19 samples took 36 minutes in a total of 3 runs, as the process is optimized during the first two days of our MFX beamtime (Table 1).

### 3.2. Determining Mpro structures with mitigated radiation damage at near physiological temperature

A total of 17 samples were screened within the scope of the study. Two of these samples contained the drug-free apo Mpro protein crystals, while the other samples contained drug-soaked protein crystals. Full datasets were collected successfully for 15 of 17 samples. Two crystal slurry samples containing native Mpro crystals soaked with montelukast and ebselen displayed high amounts of precipitation and did not show any diffraction. Analysis of the remaining 15 samples from the collected diffraction data displayed resolution ranging from 1.90 Å to 2.4 Å. In addition, the electron densities belonging to the amino acid side chains forming the catalytic cavity are well-defined (Figure 1). This region is one of the most attractive target sites for drug repurposing studies. Moreover, the critical coordinated water molecules for catalytic activity, located between His41, Cys145, His164, and Asp187 from the active site [17], were observed in 8 of 15 structures after an initial refinement process (Figure 1).

**Figure 1.**
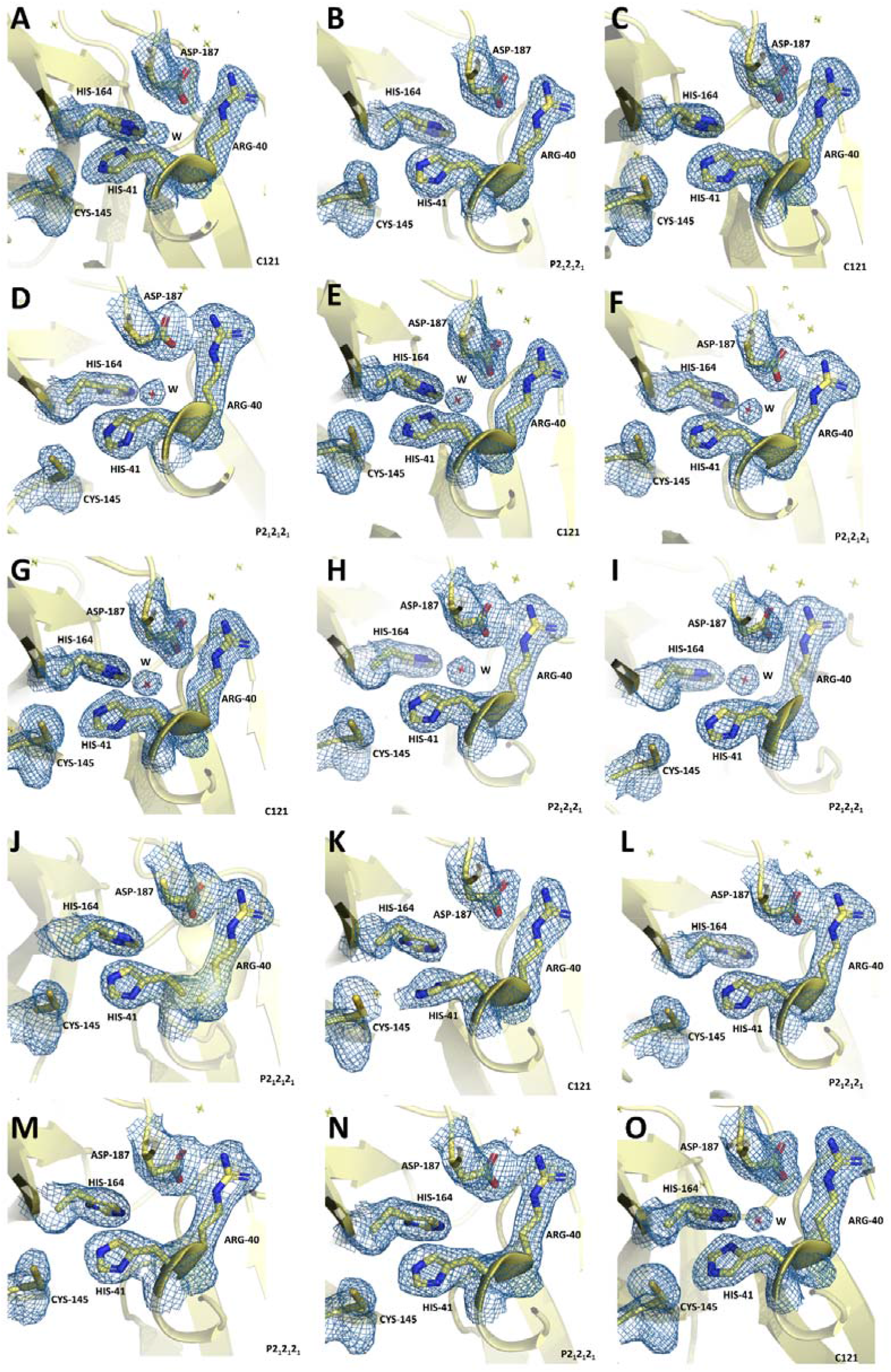
Electron density omit maps belonging to residues Arg40, His41, Cys145, His164, and Asp187 of SARS-CoV-2 Mpro structures. 2*F*o-*F*c simulated annealing-omit map is colored in slate and shown at 1.0σ level. Carbon atoms of Mpro are colored in yellow, nitrogen and oxygen atoms are colored by sky blue, and red, respectively. Space groups are shown in the bottom-right corner for each structure. Coordinated water molecules are indicated as “W”. A- Mpro9, B- Mpro19, C- Mpro13, D- Mpro14, E- Mpro 5, F- Mpro16, G- Mpro3, H- Mpro10, I- Mpro18, J- Mpro6, K- Mpro7, L- Mpro12, M- Mpro17, N −Mpro 4, O- Mpro11.

Despite these highly efficient and consistent results, target drugs could not be detected in any of the protein models after soaking at our standard 100 μg/ml concentration. Furthermore, ebselen- and montelukast-soaked crystals lost their ability to diffract in one of the two crystal forms. Precipitation in these crystal samples soaked with these two drugs may also occur as a result of binding or allosteric conformational changes. For ebselen and montelukast, cell-based and animal drug testing may provide more robust and detailed results. This is further explained in the discussion section.

### 3.3. Interpretation of experimental findings

Results are categorized under two space groups. The first space group (C121) includes six structures, while the second group (P2_1_2_1_2_1_) includes nine. The structures were aligned (Figure 2) within their respective space groups, and the RMSD values were calculated (Table 3 and Table 4). These RMSD values are indications of the similarity of refined structures, a lower RMSD indicates greater similarity. The highest RMSD value (0.54) is seen between the Mpro03 and Mpro07 structures. On the other hand, the lowest RMSD value (0.13) is observed between Mpro04 and Mpro10. All the structures in their respective space groups share the same sequence. Figure 2 shows that most conformational alterations occur at the loop region, which was expected. The helices and barrels are structurally conserved among the determined structures. The results of the data refinement process are indicated in Table 2. Ramachandran plot values in Table 2 show the percentage of residues in favorable and possible positions and outliers.

**Figure 2.**
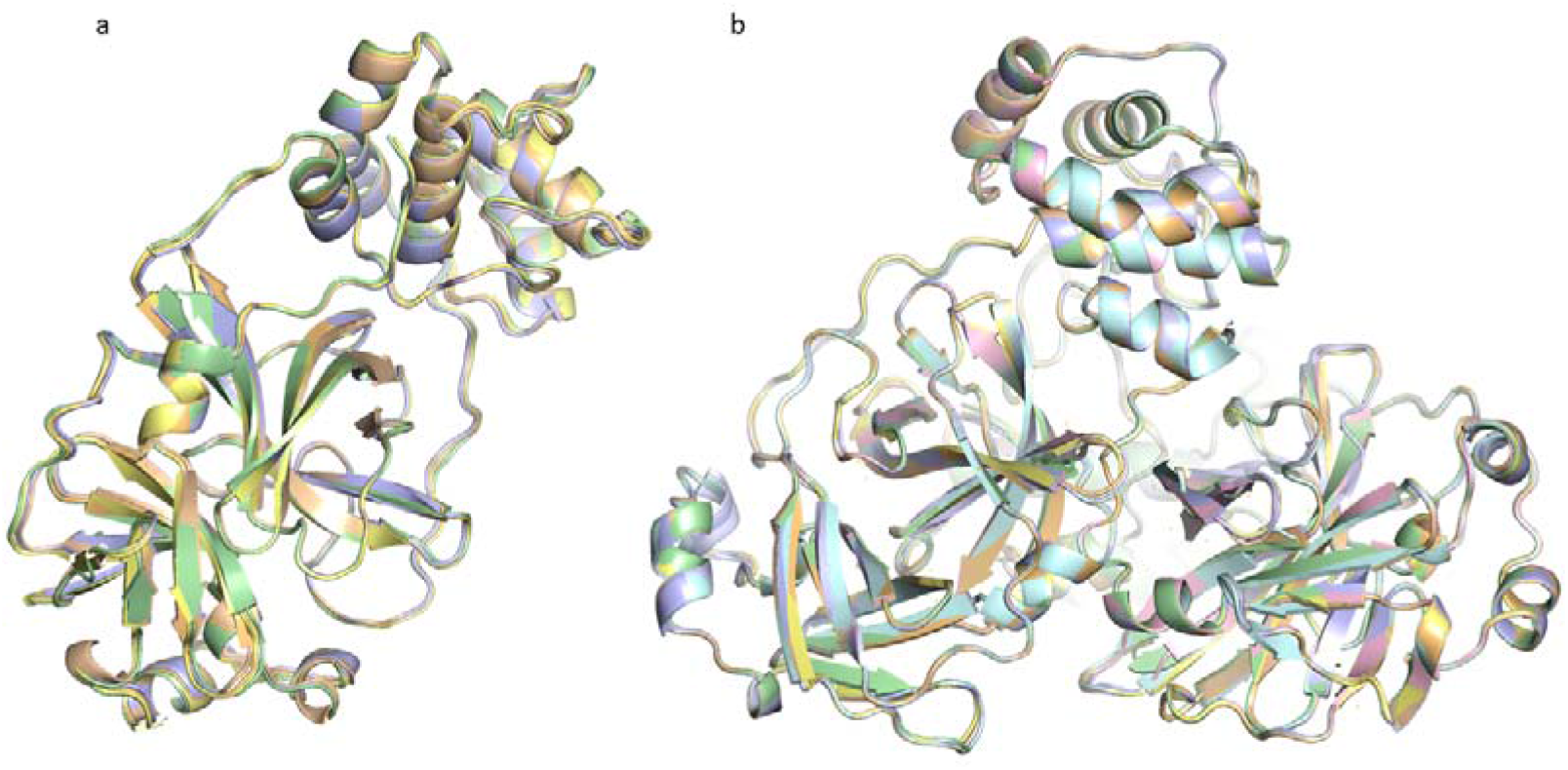
a) Structural alignment of Mpro03, Mpro07, Mpro09, Mpro11, Mpro13, all possessing similar cell constant values. Mpro03 is shown in wheat, Mpro07 in pale green, Mpro09 in light blue, Mpro11 in pale yellow, Mpro13 in light orange. RMSD values of alignments are given in Table 3 b) Structural alignment of Mpro04, Mpro06, Mpro10, Mpro12, Mpro14, Mpro17, Mpro18, Mpro19, all possessing similar cell constant values. Mpro04 is shown in wheat, Mpro06 in pale green, Mpro10 in light blue, Mpro12 in pale yellow, Mpro14 in light pink, Mpro17 in pale cyan, Mpro18 in light orange, Mpro19 in bluewhite. RMSD values of alignments are given in Table 4.

**Table 3.** RMSD values of corresponding alignments within the C121 space group.

|  | **Mpro03** | **Mpro07** | **Mpro09** | **Mpro11** |
| --- | --- | --- | --- | --- |
| **Mpro07** | 0.543 | - | - | - |
| **Mpro09** | 0.130 | 0.517 | - | - |
| **Mpro11** | 0.484 | 0.266 | 0.456 | - |
| **Mpro13** | 0.347 | 0.267 | 0.336 | 0.176 |

**Table 4.** RMSD values of corresponding alignments within the P2_1_2_1_2_1_ space group.

|  | Mpro04 | Mpro06 | Mpro10 | Mpro12 | Mpro14 | Mpro17 | Mpro18 |
| --- | --- | --- | --- | --- | --- | --- | --- |
| Mpro06 | 0.184 | - | - | - | - | - | - |
| Mpro10 | 0.132 | 0.183 | - | - | - | - | - |
| Mpro12 | 0.134 | 0.165 | 0.142 | - | - | - | - |
| Mpro14 | 0.154 | 0.138 | 0.153 | 0.154 | - | - | - |
| Mpro17 | 0.201 | 0.192 | 0.182 | 0.184 | 0.175 | - | - |
| Mpro18 | 0.160 | 0.182 | 0.144 | 0.156 | 0.145 | 0.161 | - |
| Mpro19 | 0.1180 | 0.187 | 0.172 | 0.168 | 0.158 | 0.157 | 0.137 |

## 4. Discussion

To date, the number of high-throughput data collection for drug repurposing studies conducted at XFEL is limited. There are two main reasons: the low number of XFEL facilities, and time inefficiency when compared to other light sources. Two key elements determining XFEL data acquisition rate include detector technologies and pulses per second. Thanks to the next generation detector used in this study, a high dynamic range with low-noise diffraction patterns are obtained at 120 snapshots per second, six times increase over the previous detector installed at the MFX instrument. Such advances will shorten the timeframe of high-throughput drug screenings at XFELs and make them more feasible.

The results here shown indicate that no correlation exists between effective run time and resolution. Although runtimes vary between each data collection, our final run (Mpro19) indicates that a single run can be made in as short as 36min. Getting accustomed to beamline setup and usage through online or real time data processing can help research groups use their beamtime more efficiently and allow for further high-throughput data collection.

In principle, co-crystallization and soaking are two common strategies for collecting diffraction data from protein-drug complex structures [44]. The soaking method involves ‘soaking’ the protein crystal in the drug solution at a determined concentration, whereas co-crystallization mixes the drug and the target protein prior to crystallization. Among the two techniques, co-crystallization is the least preferred because it requires additional screening time, resources, and has unpredictable crystallization success rates. The soaking method is more cost-effective and allows multiple candidate drugs to be soaked into the crystals in a short time. However, the crystal packing may not be suitable for soaking, negatively impacting ligand binding to the target site. As a result, the success rate of drug binding depends upon various crystallographic parameters and is case-specific. To overcome this limitation, we performed our soaking experiments with two different crystal forms to increase the likelihood of drug binding. Despite our attempts, we obtained structural results with only subtle differences in the active site during the HTS process. Co-crystallization, used as an alternative method, may provide more accurate structural data to demonstrate the interaction between Mpro and, not only montelukast and ebselen, but also the other drug candidates.

In this study, 9 drug candidates were screened for binding in two different crystal forms. However, none of the samples displayed noticeable experimental electron densities, suggesting at 100 μg/ml concentration soaking experiments failed. Interestingly, a high amount of precipitation was observed in two drug-protein samples containing montelukast and ebselen. Montelukast is a selective leukotriene receptor antagonist and an FDA-approved drug used in the treatment of chronic and prophylactic asthma [17]. Previously, montelukast was observed to have immunomodulatory and antiviral activities against dengue and ZIKA viruses. Therefore, it was hypothesized that montelukast may also have antiviral activity against SARS-CoV-2 [27]. Studies related to SARS-CoV-2-montelukast interaction are mostly *in silico*- [44,45,46] and *in vitro*- [17,42] oriented. Accordingly, we suggest that although montelukast interacts with Mpro, it may interfere through an allosteric effect, disrupting its conformation and perturbing the crystal lattice contacts.

In our study, we observed that ebselen-soaked P2_1_2_1_2_1_ Mpro crystals did not provide any diffraction. Similar results were demonstrated by Malla et al. [47]. Ebselen is an organoselenium molecule that displays neuroprotective, anti-microbial, and anti-inflammatory effects by forming a selenosulfide bond with the thiol groups of cysteine residues in the relevant protein [48]. Ebselen has been proven to be a potent inhibitor of SARS-CoV-2 Mpro through HTS studies [48,49]. Not only *in silico* studies [50,51] but also in crystallographic studies [52] it has been shown that ebselen can bind to the relevant region of the Mpro. Accordingly, studies on the effect of ebselen on Mpro crystals support the precipitation we observed in our Mpro-ebselen sample, and hence the potential inhibitor effect of ebselen on SARS-CoV-2. Consequently, it is likely that these two drugs interact with Mpro.

## 5. Conclusions

*De novo* drug design and discoveries require exceptionally long lead times; nevertheless, structural high-throughput data collection for drug repurposing against new emerging diseases will be one of the fastest and most effective solutions. The recent advances in XFELs and detector technologies enable structural high-throughput drug screening. In this study, we performed high-throughput SFX data collection from nine different FDA-approved drugs soaked with the two different SARS-CoV-2 main protease Mpro crystal forms. The results of our soaking experiments performed at 100 μg/ml confirmed that none of the screened drugs displayed binding to Mpro at this concentration. It is essential to perform either the soaking experiments at higher drug concentrations, or further studies with co-crystallization are required to understand the effect of candidate drugs on Mpro.

## Author Contributions

H.D. designed the experiment, H.D. performed the protein expression, purification. H.D., F.B.E., O.G. performed the crystal screening, H.D., F.B.E., O.G., O.G., C.B., E.A., S.B., G.K., Z.S., B.H., F.P., A.B., C.H.Y., C.K. performed the remote data collection. The manuscript was prepared by H.D., O.G., F.B.E., B.C., E.A., B.Y., M.G., O.G., C.B., G.U., N.T., J.J., C.H.Y.

## Data Availability Statement

The 3D electron density map of SARS-CoV-2 Main protease has been deposited in the ProteinDataBank under accession numbers 7VJW (Lopinavir, P2_1_2_1_2_1_), 7VJX (Rosuvastatin, P2_1_2_1_2_1_), 7VJY (apo form, C121), *7VJZ* (Adefovir, C121), 7VK0 (Umifenovir, P2_1_2_1_2_1_), 7VK1 (Umifenovir, C121), 7VK2 (Lopinavir, C121), 7VK3 (apo form, P2_1_2_1_2_1_), 7VK4 (Montelukast,P2_1_2_1_2_1_), 7VK5 (Adefovir,P2_1_2_1_2_1_), 7VK6 (Empagliflozin, P2_1_2_1_2_1_), 7VK7 (Atorvastatin,P2_1_2_1_2_1_), 7VK8 (Ebselen, C121). This paper does not report original code. Any additional information required to reanalyze the data reported is available from the lead contact upon request.

## Funding and Acknowledgements

The authors gratefully acknowledge use of the services and facilities of the Koç University IsBank Infectious Disease Center (KUIS-CID). HD acknowledges support from National Science Foundation (NSF) Science and Technology Centers grant NSF-1231306 (Biology with X-ray Lasers, BioXFEL). This publication has been produced benefiting from the 2232 International Fellowship for Outstanding Researchers Program of TÜBİTAK (Project Nos: 118C270, 119C132 and 120Z520). However, the entire responsibility of the publication belongs to the owner of the publication. The financial support received from TÜBİTAK does not mean that the content of the publication is approved in a scientific sense by TÜBİTAK. Use of the LCLS, SLAC National Accelerator Laboratory, is supported by the U.S. Department of Energy, Office of Science, Office of Basic Energy Sciences under Contract No. DE-AC02-76SF00515. The authors gratefully acknowledge use of the services and facilities of the Koç University Research Center for Translational Medicine (KUTTAM), funded by the Presidency of Turkey, Presidency of Strategy and Budget. The content is solely the responsibility of the authors and does not necessarily represent the official views of the Presidency of Strategy and Budget.

